# The CLAMP-Linked Invasion Protein (CLIP) plays an essential role in *Plasmodium berghei* zoites

**DOI:** 10.64898/2026.03.22.713516

**Authors:** Tanaya Unhale, Samhita Das, Carine Marinach, Sylvie Briquet, Jean-François Franetich, Lien Boeykens, Yann G.-J. Sterckx, Olivier Silvie

## Abstract

Apicomplexan parasites such as *Toxoplasma* and *Plasmodium* spp. rely on the sequential secretion of parasite apical organelles, called micronemes and rhoptries, to invade host cells. The claudin-like apicomplexan microneme protein (CLAMP) is a conserved protein that plays an essential role during host cell invasion in *Toxoplasma* and *Plasmodium* zoites. Previous studies have shown that CLAMP is essential in *Plasmodium* merozoites for erythrocyte invasion and also in sporozoites for the invasion of the mosquito vector salivary glands and of mammalian host hepatocytes. In *Toxoplasma gondii* tachyzoites, CLAMP forms a complex with two other microneme proteins, the Secreted Protein with an Altered Thrombospondin Repeat (SPATR) and the CLAMP-Linked Invasion Protein (CLIP). Both SPATR and CLIP are also expressed in *Plasmodium* sporozoites, and downregulation of SPATR impacts sporozoite infectivity in *P. berghei*. In contrast, the role of CLIP in sporozoites remains unknown. To study the function of CLIP, we used a CRISPR-assisted conditional genome editing strategy based on the dimerisable Cre recombinase in the rodent malaria model parasite *P. berghei*. Deletion of *clip* in *P. berghei* blood stages impaired parasite growth and prevented erythrocyte invasion by merozoites. Upon deletion of *clip* gene in *P. berghei* transmission stages, sporozoite development in mosquitoes was not affected, but invasion of the mosquito salivary glands was dramatically reduced. In addition, CLIP-deficient sporozoites were impaired in cell traversal and productive invasion of mammalian hepatocytes, associated with a defect in gliding motility, recapitulating the phenotype of CLAMP-deficient parasites. Collectively, our data demonstrate that CLIP plays an essential role in host cell invasion by *P. berghei* merozoites and sporozoites, and support a conserved role of the CLAMP-CLIP-SPATR complex in invasive stages of apicomplexan parasites.

## Introduction

The apicomplexan parasite *Plasmodium spp.* is the causative agent of the deadly disease malaria. The *Plasmodium* life cycle is complex and alternates between a female *Anopheles* vector and a vertebrate host. In mammals, infection begins when motile forms of the parasite known as sporozoites (SPZ) are injected in the skin during the bite of a blood-feeding infected mosquito. SPZ migrate through the dermis, invade blood vessels and are transported to the liver via the blood circulation. In the liver, SPZ migrate through several cells before infecting a final hepatocyte where the parasite differentiates into exo-erythrocytic forms (EEFs) and thousands of merozoites (MZ). Once released in the blood, MZ invade erythrocytes, initiating an exponential growth phase responsible for the malaria symptoms and complications.

Host cell invasion by *Plasmodium* and related apicomplexan parasites relies on the coordinated secretion of apical secretory vesicles (the micronemes and the rhoptries). Micronemes contain functionally diverse proteins involved in gliding motility, cell traversal and cell recognition and invasion, whereas rhoptry proteins are implicated in the formation and remodelling of a replicative niche called the parasitophorous vacuole (PV) (reviewed in [1]). In *Plasmodium* parasites, some microneme proteins are expressed in a stage-specific manner, such as the thrombospondin-related anonymous protein (TRAP) that mediates SPZ gliding motility [2], while others are expressed across invasive stages. Among the latter, the Claudin-Like Apicomplexan Microneme Protein (CLAMP) was initially identified through a genome-wide CRISPR screen in *Toxoplasma gondii* [3]. In *Toxoplasma* tachyzoites, CLAMP is secreted from the micronemes and plays an essential role during host cell invasion [3]. Depletion of CLAMP in *P. falciparum* and *P. berghei* leads to a drastic reduction of asexual blood stage growth [4,5], due to impaired erythrocyte invasion by MZ [6]. CLAMP is also expressed in SPZ of multiple *Plasmodium* species [7–9]. Conditional mutagenesis in *P. berghei* using the dimerisable Cre (DiCre) system showed that CLAMP-deficient SPZ lose their capacity to invade mosquito salivary glands and mammalian cells, associated with a defect in gliding motility [5].

In *Toxoplasma*, CLAMP forms a trimeric complex with two other microneme proteins, the Secreted Protein with Altered Thrombospondin Repeat (SPATR) and the CLAMP-Linked Invasion Protein (CLIP) [10]. Both SPATR and CLIP are predicted to be essential in *Plasmodium* blood stages [11,12], and are both expressed in SPZ [7–9]. A former study based on the Flp/FRT conditional system concluded that SPATR is required for erythrocyte (but not hepatocyte) invasion in *P. berghei* [13]. However, using another conditional knockdown strategy based on promoter swap, Costa *et al*. showed that, in the absence of SPATR, *P. berghei* SPZ fail to colonize the mosquito salivary glands, and that SPATR-deficient SPZ collected from the mosquito hemocoel are impaired in cell traversal and hepatocyte infection, associated with a strong reduction in gliding motility [14]. This phenotype is thus similar to that of CLAMP-knockout parasites, suggesting that CLAMP and SPATR could participate in the formation of a functional complex. In contrast, the role of CLIP in *Plasmodium* SPZ remains unknown.

Here, we used the DiCre conditional strategy in *P. berghei* to study the role of CLIP throughout the parasite life cycle. In the DiCre system, the Cre recombinase is split into two inactive subunits each fused to a rapamycin-ligand, which interact in the presence of rapamycin, thereby restoring Cre activity and allowing excision of Lox-flanked DNA sequences in an inducible manner [15,16]. *Plasmodium* asexual blood stage parasites can be targeted prior to transmission to mosquitoes, allowing deletion of the gene of interest and phenotypical analysis in subsequent stages of the parasite life cycle [17,18]. Conditional deletion of the *clip* gene abrogated blood stage growth and caused a dramatic decrease of mosquito salivary gland and mammalian hepatocyte invasion by SPZ. This severe phenotype was associated with a major defect in SPZ gliding motility. This study reveals that CLIP is required for *Plasmodium* SPZ motility and infectivity in both the mosquito and mammalian hosts, pointing at a possible conserved role of the CLAMP-SPATR-CLIP complex across apicomplexan parasites.

## Results

### CRISPR-mediated engineering of the *clip* gene in *P. berghei*

*P. berghei* CLIP (PBANKA_1010400) is a 277 amino acid protein with a predicted N-terminal signal peptide and C-terminal transmembrane domain. The *clip* gene comprises 4 exons and 3 introns, and is predicted to be essential in both *P. falciparum* and *P. berghei* based on genome-wide functional studies [3,10]. Previous attempts to modify the *P. falciparum clip* gene by N-terminal or C-terminal tagging were unsuccessful [10]. To study CLIP function in *P. berghei*, we used a CRISPR-assisted approach and the DiCre system, as previously described for CLAMP [6]. We genetically modified the PbCasDiCre-GFP line (which constitutively express Cas9, the DiCre components and a GFP fluorescence cassette) to introduce two LoxP sites in the *clip* gene. For this purpose, we used a one-step strategy to introduce a first LoxP site inside the first *clip* intron, and a second LoxP site immediately downstream of the STOP codon (**Figure 1A**). Using this strategy, Cre-mediated recombination results in immediate and complete suppression of CLIP expression. A triple HA tag was introduced towards the N-terminus of CLIP, just downstream of the predicted signal peptide. Two guide RNAs were selected in the *clip* locus (in the 1^st^ intron and 4^th^ exon, respectively) and cloned into a single plasmid containing PbU6 and PfU6 promoters (**Figure 1B**) [19]. The donor DNA template carrying a modified *clip* gene with two LoxP sites and a N-terminal 3xHA tag was provided as a synthetic gene and linearized before transfection. Silent mutations were introduced at the sgRNA target sites in the DNA donor template to prevent CRISPR cleavage of the recombined genomic DNA.

**Figure 1.**
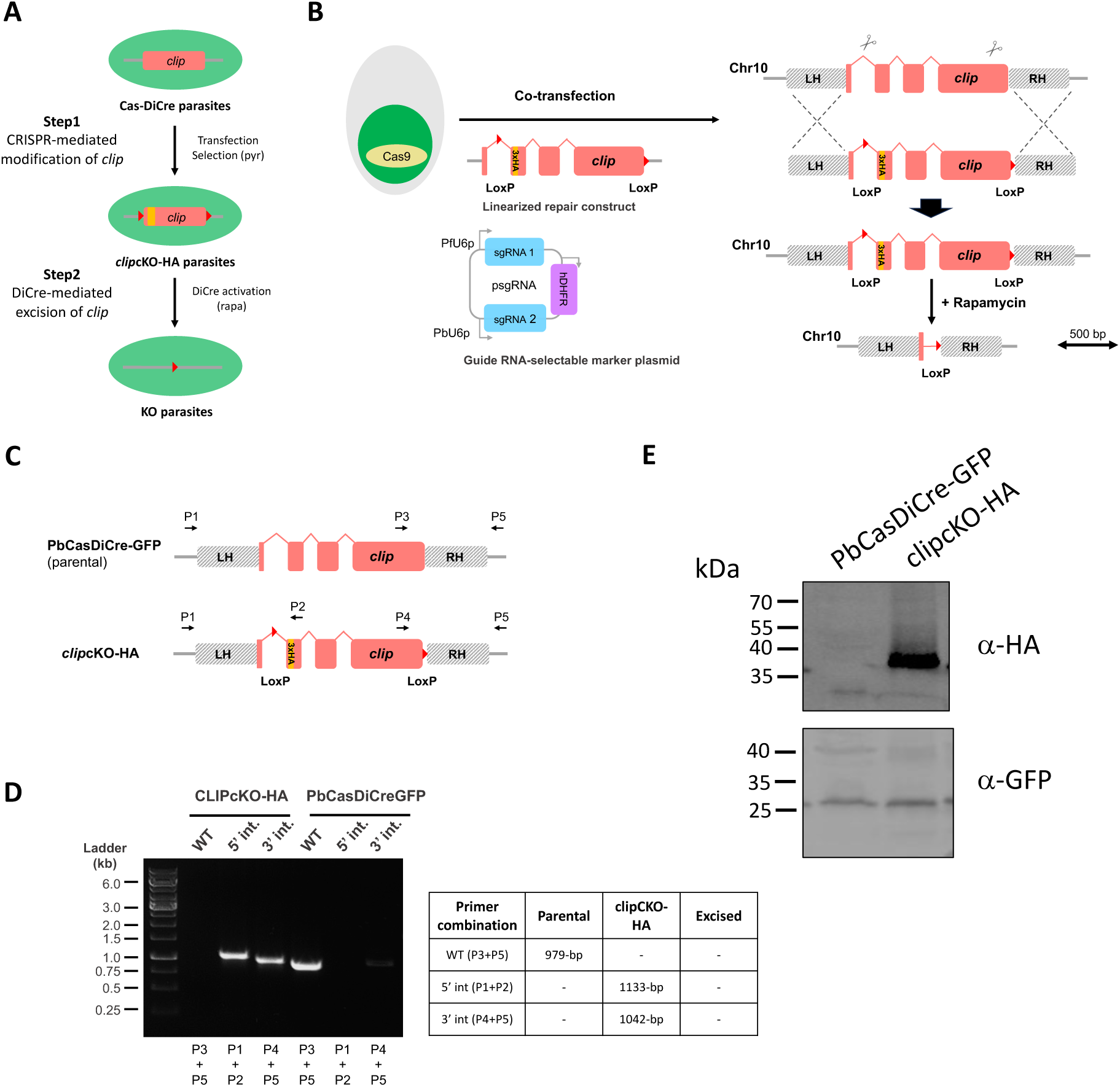
Generation of *P. berghei clip*cKO-HA parasites. **A.** Overview of the strategy used for conditional knockout of *clip* using CRISPR/Cas9 and DiCre. LoxP sites are represented as red triangles. **B.** Strategy to flox the *clip* gene in PbCasDiCre-GFP using CRISPR. PbCasDiCre-GFP parasites were co-transfected with a linearized DNA repair construct containing a floxed *clip* gene with a N-terminal 3xHA tag, and with a plasmid encoding two sgRNA guides and a pyrimethamine-resistance cassette (hDHFR). Following Cas9-mediated DNA cleavage, the repair construct was integrated by double crossover recombination at the *clip* locus, resulting in the generation of the *clip*cKO-HA parasite line. **C.** Schematic of the *clip* locus of parental and transgenic *clip*cKO-HA parasites. Genotyping primers are indicated by arrows. The loci and constructs are drawn at scale in B and C. Scale bar, 0.5 kb. **D.** PCR analysis of the genomic DNA obtained from the parental PbCasDiCre-GFP and the recombinant *clip*cKO-HA line. Confirmation of the expected recombination events was achieved with primer combinations specific for 5’ or 3’ integration. A wild type-specific PCR reaction (WT) confirmed the absence of residual parental parasites in the *clip*cKO-HA line. The size of the expected amplicons is indicated in the table. **E**. Western blot analysis of blood stage schizont lysates from *clip*cKO-HA parasites or parental PbCasDiCre-GFP parasites, using anti-HA antibodies to detect CLIP. GFP was used as a loading control.

Following the transfection of PbCasDiCre-GFP parasites with the double sgRNA plasmid targeting *clip* and the linearized donor template DNA, integration of the repair construct by double homologous recombination results in the generation of parasites containing a floxed *clip* gene with a N-terminal 3xHA tag (**Figure 1B**). Recombinant parasites were selected with pyrimethamine, and genotyped by PCR, using primer combinations specific for the parental or the recombined locus, respectively (**Figure 1C**). Parasites were then cloned by limiting dilutions and injections into mice. Genotyping PCR confirmed that the selected parasites had integrated the repair construct, with no remnant of the parental parasites in the final *clip*cKO-HA population (**Figure 1D**). PCR amplicons were verified by Sanger sequencing. Evidence of HA-tagged CLIP at the protein level was confirmed by Western blot analysis of MZ extracts, showing the presence of a ∼37 kDa band that was not detected in the parental parasite line (**Figure 1E**).

### CLIP is required for erythrocyte invasion

To assess the role of CLIP in *P. berghei* blood stages, we analyzed the effects of rapamycin on *clip*cKO parasites during blood-stage growth. Mice with patent parasitaemia (>2%) were treated with a single oral dose of rapamycin, or left untreated as a control. We then monitored the parasitaemia over time by flow cytometry to detect GFP-positive infected erythrocytes. Remarkably, rapamycin-induced excision of *clip* reduced parasitaemia to nearly zero in a single cycle (<24 hours) (**Figure 2A**), consistent with an essential role for CLIP in asexual blood stages. To verify that *clip* gene excision induced by rapamycin exposure was efficient in depleting CLIP, we exposed *clip*cKO parasites to rapamycin *in vitro* and analyzed CLIP expression by immunofluorescence using anti-HA antibodies. We observed a punctate protein distribution in untreated parasites, which was often predominant at one pole of the parasite (**Figure 2B**), reminiscent of CLAMP apical accumulation previously seen in MZ and SPZ [5,6]. In contrast, no such labelling was observed in rapamycin-treated MZ, confirming the efficiency of the conditional knockout (**Figure 2B**). We also verified the efficient depletion of CLIP by western blot analysis on MZ protein extracts (**Figure 2C**). In addition, MZ genotyping by PCR confirmed the expected DNA excision event after exposure to rapamycin (**Figure 2D**). Non-excised parasites were still detected by PCR (5’ and 3’ integration bands), and a faint excised band was detected in the absence of rapamycin exposure, suggesting some leakiness of the DiCre system.

**Figure 2.**
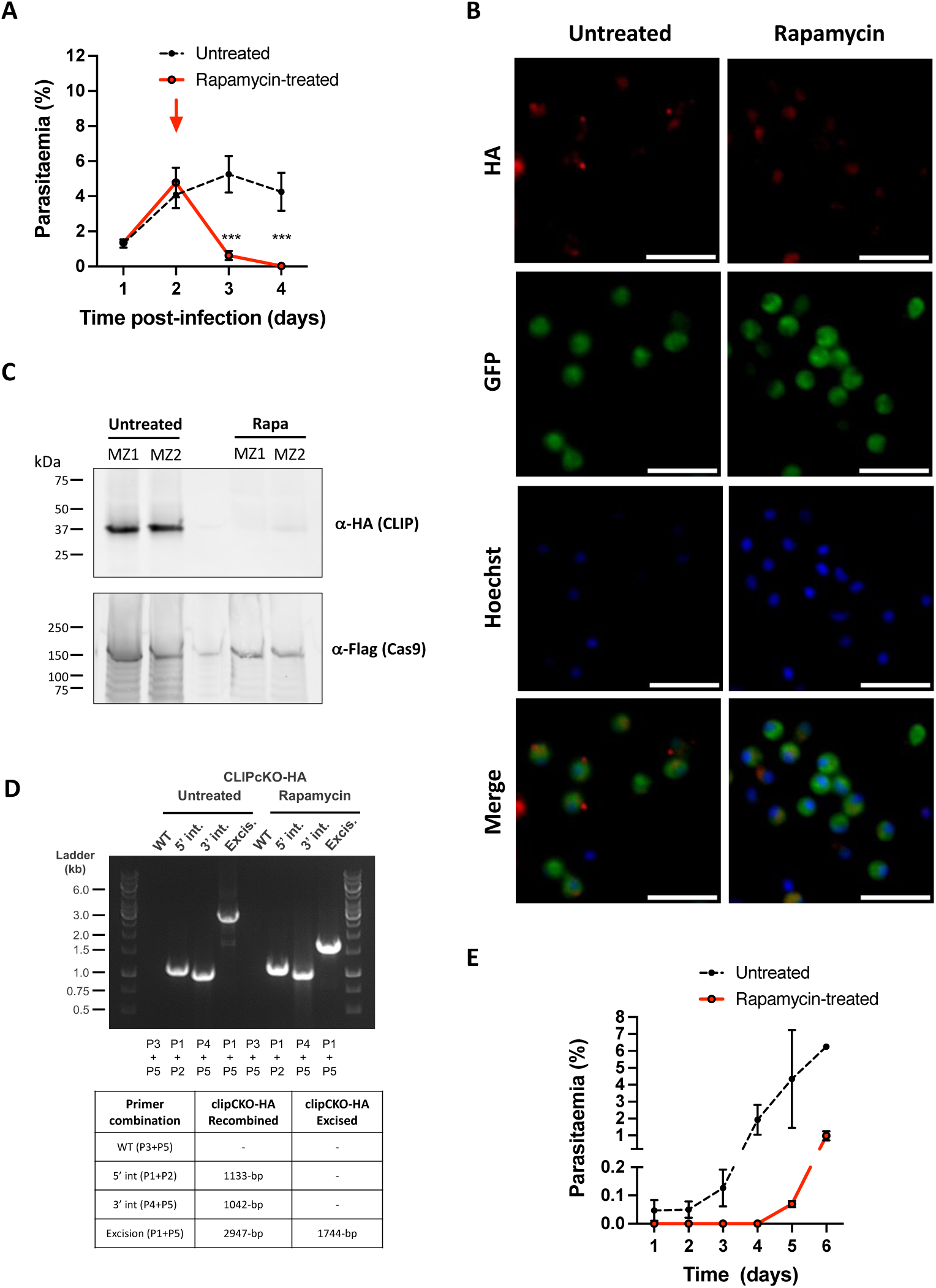
CLIP is required for *P. berghei* MZ infectivity. **A.** Blood stage growth of rapamycin-treated and untreated *clip*cKO-HA parasites. Rapamycin was administered at day 2 (red arrow). The graph shows the parasitaemia (mean +/- SEM) in groups of 4 mice, as quantified by flow cytometry based on GFP detection. ***, p<0.001 (Two-way ANOVA). **B.** Immunofluorescence analysis of *clip*cKO-HA MZ using anti-HA antibodies revealed a polar and punctate distribution of the protein in untreated parasites, and disappearance of the labelling after rapamycin exposure. Scale bar, 2 μm **C.** Western blot analysis of schizont lysates from *clip*cKO-HA parasites treated or not with rapamycin, using anti-HA antibodies to detect CLIP. Cas9-Flag was used as a loading control. Two independent samples (MZ1 and MZ2) were analyzed for each condition. **D.** PCR analysis of the genomic DNA obtained from the untreated versus rapamycin-treated cultured *clip*cKO-HA parasites, using primer combinations specific for 5’ and 3’ recombination events or gene excision. The size of the expected amplicons is indicated in the table. **E.** *clip*cKO-HA parasites were cultured for 18h in the presence or absence of rapamycin. MZ were then collected and injected intravenously into mice. The graph shows the parasitaemia (mean +/- SEM) in groups of 3 mice, as quantified by flow cytometry based on GFP detection starting 3h post-injection (day 1).

The rapid and dramatic decrease in parasitaemia during the first cycle after exposure to rapamycin suggests that CLIP-deficient parasites are unable to invade erythrocytes. To directly assess the invasive capacity of CLIP-deficient MZ, *clip*cKO MZ were produced in culture in the presence or absence of rapamycin, and injected intravenously into mice. Untreated MZ efficiently invaded mouse erythrocytes and established a blood stage infection that was patent as early as 3h post-inoculation, as evidenced by flow cytometry (**Figure 2E**). In sharp contrast, rapamycin-exposed MZ failed to invade erythrocytes, as evidenced by the absence of detectable parasitaemia after mouse inoculation (**Figure 2E**). Parasites became detectable in the blood of mice 4 days after inoculation of rapamycin-exposed parasites, consistent with the persistence of a minor population of non-excised MZ (**Figure 2D**). These data thus demonstrate that CLIP is required in *P. berghei* MZ for erythrocyte invasion.

### CLIP is required for SPZ infectivity

Next, we investigated the role of CLIP in *P. berghei* mosquito stages. For this purpose, mosquitoes were fed on *clip*cKO-infected mice, which were treated or not with rapamycin one day prior blood feeding, as previously performed for CLAMP [5]. We then monitored parasite development in the mosquito using fluorescence microscopy. Both untreated and rapamycin-exposed parasites produced oocysts, showing that CLIP is not required for parasite transmission and subsequent development in the mosquito. The salivary glands of infected mosquitoes were dissected at day 21 post-feeding and collected SPZ were counted. Rapamycin treatment of *clip*cKO parasites severely reduced the number of salivary gland SPZ **(Figure 3A)**, as previously observed with *clamp*cKO parasites [5]. In contrast, the number of hemolymph SPZ was similar in untreated (1750 SPZ/mosquito) and rapamycin-exposed *clip*cKO parasites (1500 SPZ/mosquito), indicating that CLIP is not required for SPZ development and egress from oocysts, like CLAMP [5]. We then assessed the capacity of salivary gland SPZ to traverse and invade hepatocytes *in vitro* in the absence of CLIP. We first quantified the number of traversed cells 3 hours after SPZ addition to HepG2 cell cultures by flow cytometry using a dextran-based assay as previously described [20,21]. Cell traversal was severely impaired in SPZ lacking CLIP, as shown by a dramatic reduction in the percentage of dextran-positive cells in cultures incubated with rapamycin-exposed *clip*cKO as compared to untreated SPZ **(Figure 3B)**. Host cell invasion, as determined based on the percentage of GFP-positive cells 3 hours after SPZ inoculation, was also dramatically decreased in rapamycin-treated parasites as compared to untreated controls **(Figure 3C)**. We next tested if SPZ lacking CLIP could infect and develop into EEFs in cell cultures. HepG2 cells were incubated with rapamycin-exposed or untreated *clip*cKO SPZ, followed by quantification of EEFs at 24h post-infection by flow cytometry (**Figure 3D**) or by microscopy after labelling with antibodies against UIS4 (**Figure 3E**), a marker of the PV membrane [22]. These experiments revealed an almost complete absence of EEFs in cultures inoculated with rapamycin-exposed *clip*cKO parasites **(Figure 3D-E)**. These data show that, in addition to its role during salivary gland invasion in the mosquito, CLIP is also essential in SPZ for cell traversal and productive invasion of mammalian cells. Interestingly, similar findings were observed for CLAMP [5].

**Figure 3.**
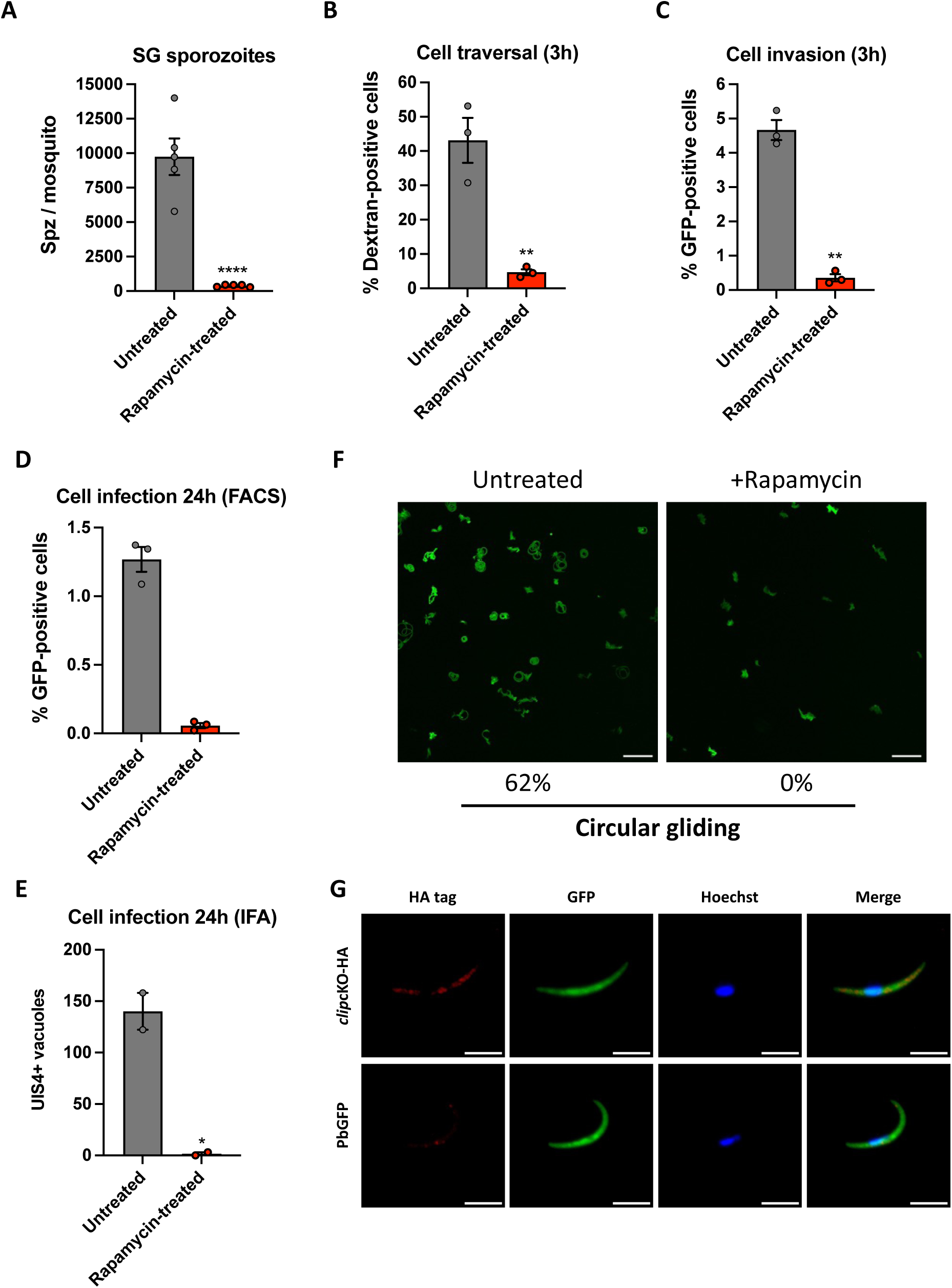
CLIP-deficient SPZ are impaired in transcellular migration, host cell infection and motility. **A.** SPZ numbers collected from salivary glands of female mosquitoes infected with rapamycin-exposed and untreated *clip*cKO parasites. The results shown are mean +/-SEM of at least three independent experiments. ****, p<0.0001 (Two-tailed ratio paired t test). **B.** Quantification of traversed (dextran-positive) HepG2 cells by FACS after incubation for 3h with rapamycin-exposed and untreated *clip*cKO salivary gland SPZ in the presence of rhodamine-labelled dextran. Results shown are mean +/- SEM of three independent experiments. **, p<0.01 (Two-tailed ratio paired t test). **C.** Quantification of invaded (GFP-positive) HepG2 cells by FACS after incubation for 3h with rapamycin-exposed and untreated *clip*cKO salivary gland SPZ. Results shown are mean +/- SEM of three independent experiments. **, p<0.01 (Two-tailed ratio paired t test). **D.** Quantification of infected (GFP-positive) HepG2 cells by FACS at 24h post-infection. Results shown are mean +/- SEM of three technical replicates from a single experiment. **E.** Quantification of UIS4-labelled EEFs in HepG2 cells as determined by fluorescence microscopy 24h post-invasion with rapamycin-exposed and untreated *clip*cKO salivary gland SPZ. Results shown are mean +/- SEM of two independent experiments. *, p<0.05 (Unpaired t test). **F**. Maximum intensity projection of video-microscopy images of untreated and rapamycin-exposed *clip*cKO salivary gland SPZ, recorded for 3 min. SPZ were activated at 37°C in the presence of serum before imaging. The proportion of SPZ displaying circular gliding was quantified in untreated (n = 79) and rapamycin-exposed (n = 35) *clip*cKO SPZ, as indicated below the images. **G.** Immunofluorescence analysis of rapamycin-exposed and untreated *clip*cKO SPZ labelled with anti-HA antibodies (red). Nuclei were stained with Hoechst 33342 (blue). Scale bar, 5 μm.

Cell traversal and productive host cell invasion both rely on the parasite gliding machinery. We hypothesized that the combined phenotype observed with parasites lacking CLIP could be caused by a defect in SPZ motility, as observed before with CLAMP [5]. Therefore, we analyzed the motility of rapamycin-exposed and untreated *clip*cKO by video-microscopy. Robust circular gliding was observed in ∼60% of untreated parasites, but not in CLIP-deficient SPZ **(Figure 3F** and **Supplementary Movies 1 and 2)**. Abrogation of motility could explain the defects observed in salivary gland invasion and in cell traversal and hepatocyte invasion, as motility is needed by the parasite to actively enter or exit cells.

Analysis of untreated *clip*cKO salivary gland SPZ by immunofluorescence using anti-HA antibodies revealed a punctate CLIP distribution in the parasite cytoplasm **(Figure 3G)**. In contrast, no labelling was observed in control SPZ of the parental line, confirming the specificity of the signal. More detailed analysis of CLIP localization using ultrastructure expansion microscopy (U-ExM) [23] showed a similar distribution of HA-tagged CLIP in SPZ as compared to TRAP, confirming that CLIP is a micronemal protein (**Figure 4A**). Occasionally, we could observe apical accumulation of CLIP at the apical tip of SPZ (**Figure 4B**), reminiscent of CLAMP distribution as seen by U-ExM [5].

**Figure 4.**
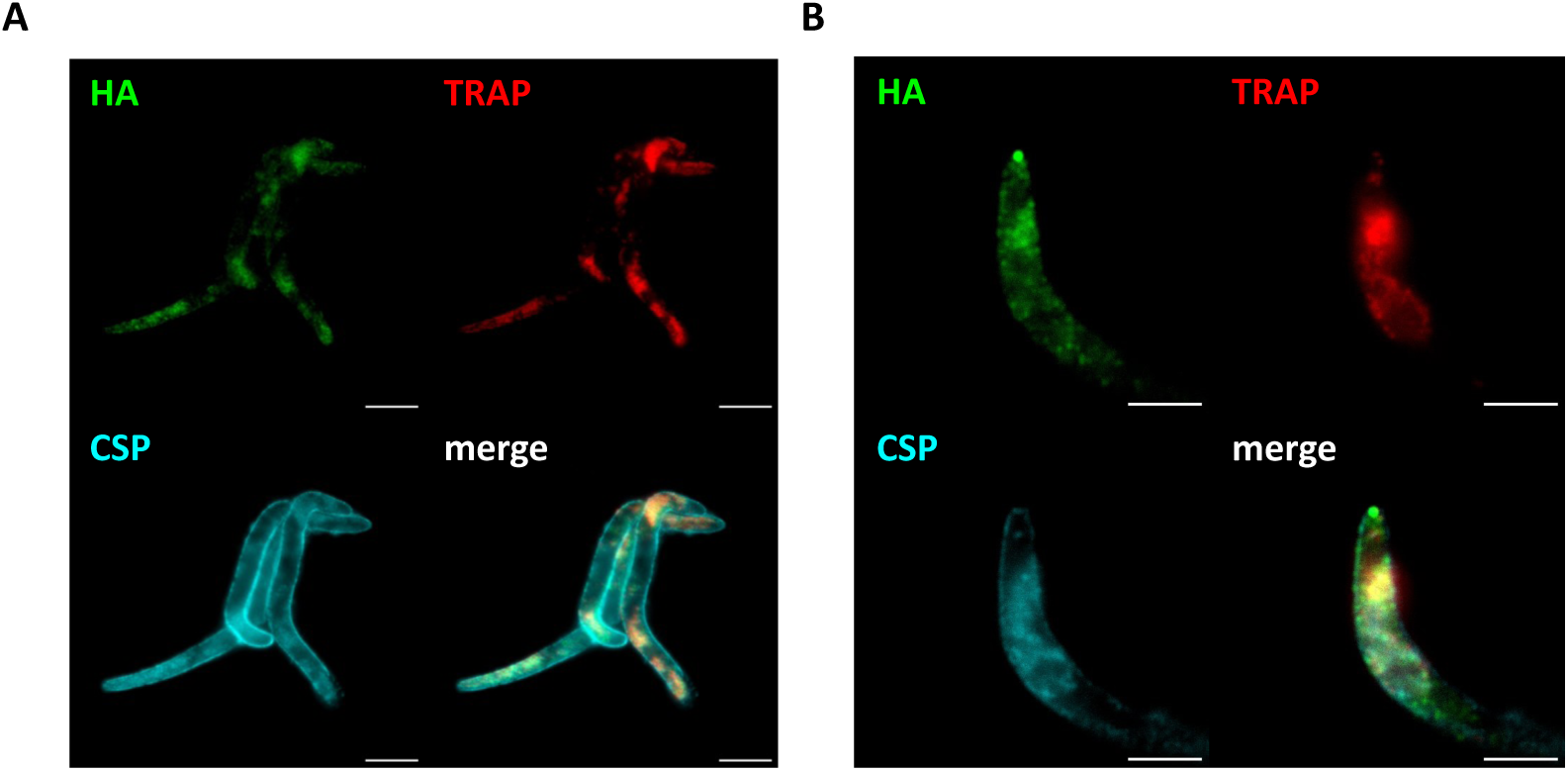
Expansion microscopy shows a micronemal distribution of CLIP in SPZ. **A.**Salivary gland SPZ expressing HA-tagged CLIP were examined by expansion microscopy using antibodies against HA (green), TRAP (red), and CSP (cyan). Scale bars, 5 μm. **B**. Occasionally, accumulation of CLIP at the apical tip of SPZ was observed. Scale bars, 2.5 μm.

## Discussion

Using a robust DiCre conditional system [6,24], we provide here a comprehensive functional characterization of CLIP in *P. berghei*. DiCre-mediated excision of *clip* abrogated CLIP expression in MZ, revealing its crucial role for erythrocyte invasion. Transmission of CLIP-deficient parasites to mosquitoes revealed that in the absence of the protein SPZ develop normally but fail to invade the salivary glands. Despite the low numbers of CLIP-deficient salivary gland SPZ, we could analyze their phenotype in functional assays. This revealed that, in addition to the defect in salivary gland invasion in the mosquito, CLIP-deficient SPZ are impaired in both traversal and invasion of mammalian hepatocytic cells. These invasion defects appear to be associated with disruption of gliding motility, given that our data demonstrate that CLIP-deficient SPZ display aberrant motility profiles.

The phenotype of CLIP-deficient mutants is similar to that previously reported with conditional mutants of *clamp* and *spatr* in *P. berghei* SPZ, with impairment of salivary gland invasion in the mosquito and hepatocyte infection in the mammalian host, associated with defects in cell traversal and gliding motility [5,6,14]. While CLAMP disruption was obtained using a similar DiCre strategy as described here with CLIP, conditional knockdown of SPATR in *P. berghei* SPZ was achieved using a promoter exchange approach [14]. Also, all three proteins appear to be indispensable in *P. berghei* blood stages [5,6,14], consistent with genome-wide screening data [25]. The similarity in the phenotypes of CLAMP, CLIP and SPATR *P. berghei* mutants strongly support the hypothesis that the three proteins form a functional complex, as reported in *Toxoplasma* [10]. This is also supported by AlphaFold-based prediction of the *P. berghei* CLAMP-SPATR-CLIP. While a high-certainty model could not be obtained for *P. berghei* CLIP alone, the prediction of the *P. berghei* CLAMP-SPATR-CLIP complex is quite robust based on local and global validation metrics (**Figure 5**). In our previous study, CLIP and SPATR were not identified by mass spectrometry after CLAMP immunoprecipitation from *P. berghei* SPZ [5]. This could reflect technical limitations and/or the transient nature of the interaction.

**Figure 5.**
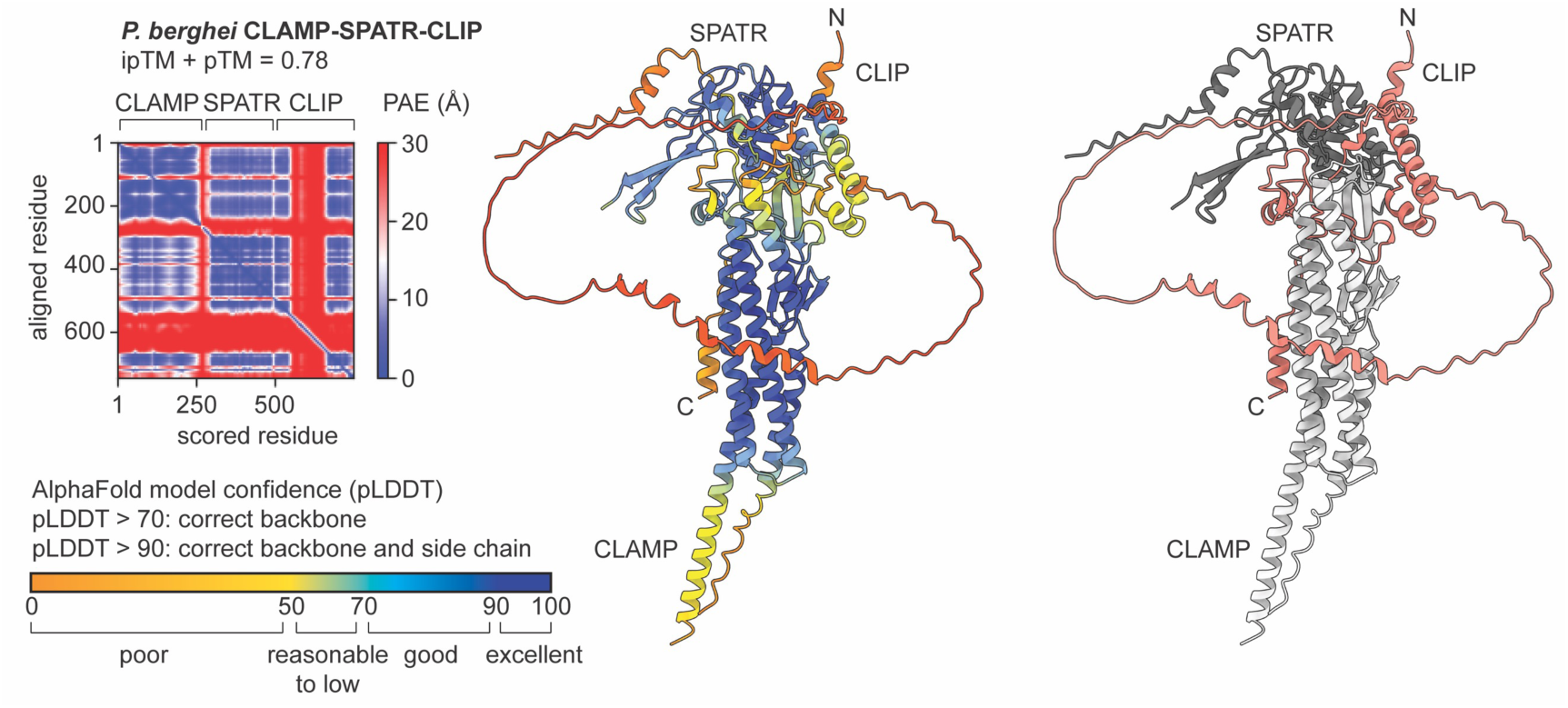
AlphaFold prediction model for the *P. berghei* CLAMP-CLIP-SPATR complex and associated quality metrics. The models are displayed as cartoon representations and are colored according to the predicted local distance difference test (pLDDT) score (middle panel), or according to the chain (right panel: CLAMP in light gray, CLIP in salmon and SPATR in dark grey). The predicted aligned error (PAE) and AlphaFold-Multimer model confidence (0.8*ipTM + 0.2*pTM) are shown on the left.

TRAP plays a pivotal role in SPZ gliding motility and was identified as a potential CLAMP-interacting partner in *P. berghei* SPZ [5]. TRAP colocalized with CLAMP in a subset of micronemes in salivary gland SPZ, and TRAP shedding was impaired in CLAMP-deficient haemolymph SPZ [5]. Whether the CLAMP-CLIP-SPATR complex participates in gliding motility via the regulation of TRAP release remains to be established. Conversely, the reduced shedding of TRAP could reflect an alteration of the protein dynamics as a consequence of disrupted motility in CLAMP-CLIP-SPATR mutants. SPATR itself could function as an adhesin in SPZ since it contains an altered thrombospondin repeat domain [14]. Our U-ExM data support a micronemal distribution of CLIP in *P. berghei* SPZ, as reported for CLAMP [5]. Also, similarly to CLAMP [6], CLIP shows a punctate distribution in MZ, presumably at the apex of the parasite, although the nature of this compartment remains to be established.

Interestingly, SPATR is essential in *Plasmodium* but dispensable in *Toxoplasma* [26–28], and, like CLAMP, is required for gliding motility in *Plasmodium* SPZ but not *Toxoplasma* tachyzoites [4,26–28]. The phenotypical differences observed with CLAMP and SPATR mutants in *Plasmodium* SPZ versus *Toxoplasma* tachyzoites could reflect species-specific functional diversification or stage-specific functions of the CLAMP-CLIP-SPATR complex, considering the highly migratory behavior of *Plasmodium* SPZ. As noted previously for CLAMP, it is also plausible that the complex has two functions, one in regulating microneme functions (such as gliding motility) in SPZ, and rhoptry functions (host cell invasion) in *Plasmodium* MZ (and possibly SPZ) and *Toxoplasma* tachyzoites. Structure function analysis showed that the C-terminal proline rich domain of CLAMP plays an essential role in *Toxoplasma*, raising the possibility that this part of the protein is involved in signal transduction required for rhoptry discharge [10]. The same authors proposed that the CLAMP-CLIP-SPATR complex mediates rhoptry discharge following host cell contact. However, a more recent study from the same group identified another trimeric complex of microneme proteins (MIC1/4/6) as being directly involved in triggering rhoptry discharge in *Toxoplasma*, via recognition of N-glycans on the host cell membrane [29]. Whether the CLAMP-CLIP-SPATR complex acts upstream or downstream of host cell recognition thus remains to be determined.

In summary, we show here that CLIP plays an essential role in *P. berghei* MZ and SPZ, supporting a conserved role of the CLAMP-CLIP-SPATR complex in apicomplexan zoites. This complex may thus represent a potential new target for antimalarial strategies. Previous reports showed that antibodies against SPATR can block invasion of hepatoma cells by *P. falciparum* SPZ [30], and invasion of human erythrocytes by *P. knowlesi* MZ [31]. Also, CLAMP-specific antibodies could inhibit invasion of equine erythrocytes by the apicomplexan *Theileria equi*, by targeting neutralization-sensitive epitopes present in the predicted extracellular loops of the protein [32]. Targeting the CLAMP-CLIP-SPATR complex could thus open interesting perspectives for interventions acting both against both MZ (to prevent invasion of erythrocytes) and SPZ (to block infection early after transmission by the mosquito).

## Materials and methods

### Ethics Statement

All animal work was conducted in strict accordance with the Directive 2010/63/EU of the European Parliament and Council on the protection of animals used for scientific purposes. Protocols were approved by the Ethics Committee Charles Darwin N°005 (approval #45554-2023102311026657). The study was conducted in accordance with ARRIVE guidelines (https://arriveguidelines.org).

### Experimental animals, parasites and cell lines

Female Swiss mice (4–8 weeks old, from Janvier Labs) were used for all routine parasite infections. Parasite transfections were performed in the parental PbCasDiCre-GFP line (ANKA strain), which constitutively express Cas9 and the DiCre components in addition to a GFP fluorescence cassette [6]. Parasite infections in mice were initiated through intraperitoneal injections of infected RBCs. A drop of blood was collected from the tail in 1ml PBS daily and used to monitor the parasitaemia by flow cytometry. *Anopheles stephensi* mosquitoes were reared at 24-26°C with 80 % humidity and permitted to feed on anaesthetised infected mice, using standard methods of mosquito infection as previously described [33]. Post-feeding, *P. berghei*-infected mosquitoes were kept at 21°C and fed on a 10% sucrose solution. Salivary gland SPZ were collected at day 21 post-infection by hand dissection and homogenisation of isolated salivary glands in complete DMEM medium (supplemented with 10% FBS, 1% Penicillin-Streptomycin and 1% L-Glutamine), then counted in a Neubauer haemocytometer. HepG2 cells (ATCC HB-8065) were cultured in complete DMEM in collagen-coated plates, at 37°C, 5% CO2, as previously described [34], and used for invasion and cell traversal assays.

### DNA constructs

#### sgRNA guide plasmids

Using the Chop-Chop (https://chopchop.cbu.uib.no) and Benchling (https://www.benchling.com) programs, two 19-20 bp guide RNA sequences were selected upstream of PAM motifs in the 5’ and 3’ regions of *clip* gene, and used for the design of complementary oligonucleotides. A guanosine nucleotide was added at the 5’ end of the forward oligonucleotide for enhancing transcriptional initiation [19]. Paired oligonucleotides were annealed and sequentially cloned into the psgRNA_Pf-Pb U6_2targets plasmid at the *Bsm*BI and *Bsa*I sites, using the In-Fusion HD Cloning Kit (Takara), resulting in the insertion of the guide RNA immediately downstream of PfU6 or PbU6 promoter, respectively [19]. This plasmid contains a *hDHFR*-*yfcu* cassette, for positive selection by pyrimethamine and negative selection by 5-fluorocytosine (5-FC) [35,36]. The resulting sgRNA guide plasmid was checked by Sanger sequencing prior to transfection.

#### Donor DNA templates for DNA repair by double homologous recombination

The donor DNA template to modify the *clip* locus was provided as a synthetic gene, containing a 500-bp 5’ upstream fragment immediately upstream of the ATG of *clip*, serving for 5’ homologous recombination, a 40-bp sequence corresponding to the first exon, a 186-bp fragment corresponding to a modified first intron that includes a LoxP site and mutations at the sgRNA target site, a 126-bp fragment corresponding to the second exon containing an inserted 3xHA epitope tag, a 380-bp fragment corresponding to introns 2 and 3 and exon 3, a 578-bp fragment corresponding to a modified exon 4 with mutations introduced at and downstream of the sgRNA target site, followed by a second 34-bp LoxP site and a 500-bp 3’ downstream fragment serving for 3’ homologous recombination. The entire synthetic gene was flanked by two *Xho*I sites allowing linearization prior to transfection. All the DNA sequences are listed in **Supplementary Table S1**.

### Parasite transfection

For parasite transfection, schizonts were purified from an overnight culture of the parent parasite line PbCasDiCre-GFP [6] and transfected with a mix of 10 μg of sgRNA plasmid and 10 μg of linearized donor template by electroporation using the AMAXA Nucleofector device (program U033), as previously described [37], and immediately injected intravenously into the tail vein of SWISS mice. To permit the selection of resistant transgenic parasites, pyrimethamine (35 mg/L) was added to the drinking water and administered to mice, starting one day after transfection and for a total of 4-5 days. Following withdrawal of pyrimethamine, the mice were monitored daily by flow cytometry to detect the reappearance of parasites. When parasitaemia reached at least 1%, mice were anaesthetized by exposure to 3.5% isoflurane and euthanized under anaesthesia via terminal blood draw followed by cervical dislocation. Mouse blood was collected for preparation of cryostocks and isolation of parasites for genomic DNA extraction and genotyping. Parasites were cloned by limiting dilution and injection into mice to generate the final *clip*cKO-HA population used in this study.

### Genotyping PCR

The blood collected from infected mice was passed through a CF11 column (Whatman) to deplete leucocytes. The collected RBCs were then centrifuged and lysed with 0.2% saponin (Sigma), before genomic DNA isolation using the DNA Easy Blood and Tissue Kit (Qiagen), according to the manufacturer’s instructions. Specific PCR primers were designed to check for wild-type and recombined loci and are listed in **Supplementary Table S1**. PCR reactions were carried out using Recombinant *Taq* DNA Polymerase (Thermo Scientific) and standard PCR cycling conditions.

### Rapamycin-induced gene excision

*clip*cKO-infected mice were administered a single dose of 200 μg rapamycin (Rapamune, Pfizer) by oral gavage. Rapamycin-treated *clip*cKO parasites were transmitted to mosquitoes one day after rapamycin administration to mice, as described [17,38]. Excision of *clip* ex vivo in blood stages was achieved by culturing infected erythrocytes for 18 hours in the presence of 100 nM rapamycin (from 1 mM stock solution in DMSO, Sigma-Aldrich). Control cultures were exposed to DMSO alone (0.01% vol/vol).

### *In vitro* infection assays

HepG2 cells cultured in DMEM complete medium were seeded at a density of 30,000 cells/well in a 96-well plate for flow cytometry analysis or 100,000 cells/well in 96-well µ-slide (Ibidi) for immunofluorescence assays, 24 hours prior to addition of SPZ. Culture medium was refreshed with complete DMEM on the day of infection, followed by incubation with 10,000 or 1,000 SPZ, for flow cytometry or immunofluorescence assays, respectively. For quantification of traversal events, rhodamine-conjugated dextran (0.5 mg/ml, Life Technologies) was added to the wells together with SPZ. After 3 hours, cells were washed twice with PBS, trypsinized, then resuspended in complete DMEM for analysis by flow cytometry on a Guava EasyCyte 6/2L bench cytometer equipped with 488nm and 532nm lasers (Millipore). Control wells were prepared without SPZ to measure the basal level of dextran uptake. To quantify parasite liver stage infection, cells were washed twice with complete DMEM 3 hours after SPZ addition, and then incubated for another 24-36h. Cultures were then either trypsinized for quantification of GFP-positive cells by flow cytometry, or fixed with 4% PFA for immunofluorescence analysis. Following two washes with PBS, cells were then quenched with 0.1M glycine for 5 min, washed twice with PBS then permeabilized with 1% Triton X-100 for 5 min before 2 washes in PBS and blocking in PBS + 3% BSA. Samples were then stained with goat anti-UIS4 primary antibodies (1:500, Sicgen), followed by donkey anti-goat Alexa Fluor 594 secondary antibodies (1:1000, Life Technologies), both diluted in PBS with 3% BSA. EEFs were then counted based on the presence of a UIS4-stained PV.

### Motility assay

For motility assays, rapamycin-exposed and untreated *clip*cKO SPZ were collected by manual dissection of infected mosquitoes and kept in PBS on ice until imaging in RPMI medium supplemented with 10% foetal calf serum (Life Technologies). SPZ were then deposited in an 8-well chamber slide (Ibidi), centrifuged for 6 min at 100 x g, 4°C, and placed at 37°C in the microscope chamber with 5% CO_2_. Acquisitions were initiated after 3 min, to allow SPZ activation, with one image captured every second for a total of 3 min. Stacks of 9 images with a 1,5 µm z step size were acquired on a Nikon CREST V3 spinning disk microscope equipped with a 40x/NA:1,15 water immersion objective and a Photometrics Kinetix sCMOS cameras, and further processed in a z-projection of max intensities with NIS-Element acquisition software. All the generated z-projection images (181 images corresponding to 1 image per second for 3 minutes) were superimposed to analyze SPZ motility.

### Western-blot analysis

Parasites were resuspended in Laemmli buffer and analyzed by SDS-PAGE under reducing conditions. Western blotting was performed using a rat monoclonal antibody against HA (3F10, Roche), a mouse monoclonal against Flag (M2, Sigma) or goat polyclonal antibodies against GFP (Sicgen), and secondary antibodies coupled with Alexa Fluor 680 or 800. Membranes were then analyzed using the InfraRed Odyssey system (Licor).

### Immunofluorescence assays

SPZ were collected by disruption of salivary glands from mosquitoes infected with *clip*cKO-HA or parental parasites, and resuspended in PBS, placed on a coverslip then fixed for 10 min with 4% PFA followed by two washes with PBS. For immunostaining, SPZ were permeabilized with Triton X-100 and incubated with 3% BSA in PBS for 1h. SPZ were then stained with anti-HA primary rat antibody (3F10, Roche), washed twice with PBS then incubated with Alexa Fluor 594-conjugated goat anti-rat secondary antibodies (Life Technologies) and Hoechst 33342 (Life Technologies), before examination by fluorescence microscopy. For immunofluorescence on infected erythrocytes, cultures of untreated or rapamycin-treated *clip*cKO-HA schizonts were fixed in 4% PFA as described above and permeabilized with Triton X-100. The HA tag was revealed using a rat monoclonal antibody (3F10, Roche) and Alexa Fluor 594-conjugated goat anti-rat secondary antibodies (1:1000, Life Technologies). Nuclei were stained with Hoechst 33342 (Life Technologies). All images were acquired on an Axio Observer Z1 Zeiss fluorescence microscope using the Zen software (Zeiss). The same exposure conditions were maintained for all the conditions in order to allow comparisons. Images were processed with ImageJ for adjustment of contrast.

### Ultrastructure expansion microscopy

SPZs were collected from infected mosquito salivary glands, centrifuged at 3800 g during 4 min at 4 °C and resuspended in 1X PBS. Parasites were sedimented on poly-D-lysine coverslips (100 μL/coverslip) during 30 min at room temperature (RT). Samples were then prepared for U-ExM as previously described [5]. Briefly, coverslips were incubated overnight in a 2% Acrylamide/1.4% Formaldehyde solution at 37 °C. Gelation was then performed in 10% ammonium persulfate (APS)/10% Temed in monomer solution (19% Sodium Acrylate; 10% Acrylamide; 0.1% BIS-Acrylamide in PBS) during 1 h at 37 °C. Following gelation, denaturation was performed in 200mM SDS, 200mM NaCl and 50mM Tris pH 9.0 during 90 min at 95 °C. A first round of expansion was performed by incubating the gels thrice in ultrapure water for 30 min at RT. Gels were then washed in PBS twice for 15 min to remove excess water and blocked with 2% BSA in PBS for 30 min at RT. Staining was then performed by incubation with primary antibodies diluted in PBS containing 2% BSA at RT overnight with 120–160 rpm shaking. We used antibodies against HA (3F10, Roche), TRAP [39] and CSP (MRA-100A, BEI Resources). The next day, gels were washed 3 times for 10 min in PBS-Tween 0.1%. Incubation with the secondary antibodies was performed for 3 h at RT with 120–160 rpm shaking, followed by 3 washes of 10 minutes in PBS-Tween 0.1%. Directly after washing, gels were expanded for a second round in ultrapure water for 30 min, thrice. For imaging, 5 mm x 5 mm gel pieces were cut from the expanded gels and mounted between glass slides and Poly-D-Lysine coated coverslips. Acquisitions were made on a Zeiss Axio Imager Z1 fluorescence microscope equipped with a Plan-Apochromat 100x/1.40 Oil DIC M27 objective, using the AxioVision software (Zeiss). Images were processed with ImageJ for adjustment of contrast.

### AlphaFold-based structure prediction

The structural model of the *P. berghei* CLAMP-SPATR-CLIP complex was predicted using AlphaFold-Multimer [40]. Twenty-five models were predicted per run and the best models underwent a final relaxation step. The models were evaluated based on the following parameters: AlphaFold-Multimer model confidence (a weighted combination of the predicted template modeling (pTM) and interface predicted template modeling (ipTM) scores, 0.8*ipTM + 0.2*pTM) [40], the predicted aligned error (PAE) matrix [41–43], and the local and global predicted local distance difference test (pLDDT) scores [41–43]. Molecular graphics and analyses were performed with UCSF ChimeraX [44].

### Statistical Analysis

Statistical significance was assessed by two-way ANOVA, Log-rank (Mantel-Cox) or ratio paired t tests, as indicated in the figure legends. All statistical tests were computed with GraphPad Prism 10 (GraphPad Software). *In vitro* experiments were performed with a minimum of three technical replicates per experiment. Quantitative source data are provided in **Supplementary Table S2**.

## Acknowledgements

We thank Thierry Houpert and Louise Deltour Foglio for rearing of mosquitoes. This work was funded by grants from the Laboratoire d’Excellence ParaFrap (ANR-11-LABX-0024) and the Agence Nationale de la Recherche (ANR-20-CE18-0013), and was supported by the Conseil Régional d’Île-de-France via the DIMOneHealth action for funding of the spinning disk confocal microscope. The authors acknowledge use of the CalcUA and VSC supercomputing facilities and wish to thank the staff for outstanding support.

## Supporting information

### Supplemental tables

**S1 Table**. Sequence of oligonucleotides and synthetic genes used in the study.

**S2 Table**. Quantitative source data and statistical analysis.

### Supplemental movies

**Movie 1. Motility of untreated *clip*cKO salivary gland SPZ**. Motility was recorded with one frame per second (fps) for 3 min after activation at 37°C. Video was edited with 7fps.

**Movie 2. Motility of rapamycin-treated *clip*cKO salivary gland SPZ**. Motility was recorded with one frame per second (fps) for 3 min after activation at 37°C. Video was edited with 7fps.

